# A subset of type 4 secretion system effectors of *Brucella* spp. associates to outer membrane vesicles

**DOI:** 10.1101/2025.10.06.680763

**Authors:** Maren Ketterer, Naadir Ganief, Petra Chiquet, Adélie Lannoy, Audrey Verhaeghe, Joanne Braunweiler, Marc Dieu, Xavier De Bolle, Maxime Québatte, Christoph Dehio

**Affiliations:** Biozentrum, University of Basel, Basel, Switzerland; URBM and Department of Biology, Namur Research Institute for Life Sciences (NARILIS), Universtiy of Namur, Namur, Belgium; Laboratoire de Microbiologie et Génétique Moléculaires, Centre de Biologie Intégrative, Centre National de la Recherche Scientifique, Université de Toulouse, UPS, Toulouse, France; MaSUN, Mass Spectrometry Facility,University of Namur, Namur, Belgium; WEL Research Institute avenue Pasteur, 6 1300 Wavre, Belgium

**Keywords:** *Brucella*, OMV, effectors, T4SS, translocation, secretion

## Abstract

The establishment of a replicative niche in the hostile environment of the host presents an enormous challenge for pathogens. Intracellular pathogens such as *Brucella* spp., the Gram-negative causative agents of Brucellosis, must subvert diverse host functions to ensure survival and replication. One of the key adaptations to achieve this is the translocation of effector proteins into host cells via its type 4 secretion system (T4SS), a key virulence factor. But effector identification in *Brucella* is particularly challenging, as previously identified effectors lack a conserved translocation signal, exhibit variable requirements for translocation, and in some cases appear to be translocated in a T4SS-independent manner.

Here, we demonstrate that a subset of previously described T4SS effector proteins associates with outer membrane vesicles (OMVs) in different *Brucella* species. Most of these effector proteins encode predicted signal peptides for periplasmic export or transmembrane domains. Among them, BspC and VceA carry functional signal peptides that direct their export into the periplasm in a Sec-dependent manner. From the periplasm these proteins are subsequently secreted into the extracellular milieu, likely via the formation and release of OMVs. Our findings provide new insights into protein secretion by *Brucella*, suggesting that OMVs may represent an alternative secretion pathway to the T4SS.

## Introduction

*Brucella* spp. are Gram-negative, facultative intracellular bacteria and the causative agents of the zoonosis Brucellosis. The genus is divided in classical species (e.g., *Brucella melitensis*, *Brucella abortus*, *Brucella suis*) and “atypical” species (e.g., *Brucella microti*), distinguished by their bacteriological features, host specificity, and pathogenic potential (reviewed in (Occhialini *et al*. 2022)). To ensure survival and replication *Brucella* manipulates host cell functions through its T4SS, which is essential for these processes. Following uptake into host cells, *Brucella* initially follows the canonical endocytic pathway (Pizarro-Cerdá *et al*. 1998a, 1998b; Comerci *et al*. 2001; Celli *et al*. 2003; Bellaire, Roop and Cardelli 2005), where transient acidification of the late endosome is required for T4SS expression (Porte, Liautard and Köhler 1999; Boschiroli *et al*. 2002; Altamirano-Silva *et al*. 2018), and effector proteins are thought to mediate conversion of the non-replicative *Brucella* containing vacuole (BCV) to the replicative BCV, which is directly connected to the ER (Pizarro-Cerdá *et al*. 1998b; Lestrate *et al*. 2000; Comerci *et al*. 2001; Celli *et al*. 2003; Bellaire, Roop and Cardelli 2005; Starr *et al*. 2008; Sedzicki *et al*. 2018).

Interestingly, while the *virB* operon encoding the T4SS is indispensable for intracellular replication and essential in a transposon mutant screen, deletion of individual effector genes rarely reduces virulence, suggesting redundancy (Myeni *et al*. 2013; Marchesini *et al*. 2016; Sternon *et al*., 2018, Smith *et al*. 2020; Li *et al*. 2022).

Moreover, the mechanisms governing effector recognition and translocation remain poorly understood. *Brucella* lacks the classical VirD4 coupling protein, which is essential for effector recognition in related T4SSs (O’Callaghan *et al*. 1999; Schulein and Dehio 2002; Schulein *et al*. 2005). Further, *Brucella* effectors do not share a consensus translocation signal designating them as T4SS substrates as described in related systems (Huang *et al*. 2011; Stanger *et al*. 2017; Wagner, Tittes and Dehio 2019). Instead, translocation requirements vary between effectors and may involve the N-terminus (e.g., BPE123 (Marchesini *et al*. 2011)), the C-terminus (e.g., VceC (De Jong *et al*. 2008)), or both (e.g., BspA (Myeni *et al*. 2013)). Other effectors (e.g., BspH, BspJ, BPE865) appear to be translocated independently of the T4SS when N-terminally tagged, yet require the presence of the T4SS when C-terminally tagged (Myeni *et al*. 2013). And some effectore (e.g. BspG) are entirely T4SS-independent (Myeni *et al*. 2013). Collectively, these observations suggest that *Brucella*’s T4SS exhibits unusual flexibility in substrate recognition and may operate in concert with additional, as yet undefined, secretion or translocation pathways that might depend on the T4SS-driven subcellular localization of the bacterium. However, *Brucella* lacks other classical secretion systems such as the type 2, 3, 9 secretion systems, and the T4SS is regarded as the principal pathway for effector delivery.

However, several species produce outer membrane vesicles (OMVs) (Avila-Calderón *et al*. 2012, 2020; Pollak *et al*. 2012; Araiza-Villanueva *et al*. 2019; Socorro Ruiz-Palma *et al*. 2021) - nanostructures ubiquitously shed by Gram-negative bacteria, composed of outer membrane and periplasmic components. These *Brucella*-derived OMVs can modulate immune responses and promote bacterial internalization, but their molecular cargo and role in pathogenicity and virulence remain poorly understood (Pollak *et al*. 2012; Avila-Calderón *et al*. 2020).

Here, we investigated the protein content of OMVs from different *Brucella* species to determine whether described effector proteins localize to these vesicles. Using comparative proteomics we identified a subset of effectors which are consistently present in OMVs of different species, many of which contain predicted signal peptides for periplasmic targeting. Focusing on BspC and VceA, we show that both carry functional signal peptides and are therefore exported to the periplasm. They are further secreted to culture supernatant in a T4SS-independent manner, and localize to an insoluble, OMV-containing fraction. These findings suggest that *Brucella* may use OMVs as an alternative seretion pathway and highlight the need to explore non-canonical secretion mechanisms in this pathogen.

## Material and Methods

### Bacterial strains and growth conditions

Bacterial strains used in this study are listed in Table S1. *E. coli* strains were cultivated in lysogeny broth (LB) or on Luria-Bertini agar (LA) plates at 37°C overnight supplemented with appropriate antibiotics.

*Brucella abortus* 2308 and *Brucella microti* CCM4915T strains were grown in tryptic soy broth (TSB, Sigma-Aldrich 22092) or defined medium (DM) (De Barsy *et al*. 2011) at 37°C with agitation supplemented with appropriate antibiotics. Solid cultures were grown on LA agar or tryptic soy agar (TSA, Difco 236950) plates with appropriate antibiotics at 37°C. Cultures were inoculated directly from frozen stocks stored at −80°C.

### Cloning

All manipulations with DNA were performed following standard techniques, and plasmid integrity was verified by sequencing. Plasmids and primers are listed in Table S2 and S3, respectively. Plasmids were conjugated into the different *Brucella* species following standard protocols.

### Deletion of virB9 in Brucella microti

*virB9* was deleted from *B. microti* in frame by double homologous recombination using the SacB allelic exchange suicide vector pNPTS138, which confers kanamycin resistance and sucrose sensitivity upon insertion. Excision of the insertion was counterselected on 5% sucrose TSA plates. For further details see (Deghelt *et al*. 2014). Correct deletion was confirmed by colony PCR using primers listed inTable S3 and via sequencing.

### Alkaline phosphatase assay

*B. microti* strains expressing the different alkaline phosphatase fusions were grown for 6 h in TSB at 37°C with 170 rpm in presence of 100 ng/ml anhydrotetracycline (aTc) to induce protein expression. Volumes equal to an OD_600nm_ of 2 in 1 ml were spun down, resuspended in 100 µl fresh TSB, 200 μg/ml 5-Bromo-4-Chloro-3-Indolyl Phosphate (BCIP, Sigma-Aldrich, B6149) was added, and samples were incubated at 37°C for 1-2 h before imaging.

### Axenic secretion assay

Overnight cultures were grown in 2YT in the presence of 34 μg/ml chloramphenicol until stationary phase. The secretion culture was inoculated to an OD_600nm_ of 0.4 in 10 ml defined medium (DM) pH 7, supplemented with 17 μg/ml chloramphenicol, as higher chloramphenicol concentrations inhibited growth in DM. Secretion cultures were incubated at 37 °C at 170 rpm for 7.5 h for *B. microti* and 17 h for *B. abortus* with 100 ng/ml aTc to induce expression of fusion proteins. Samples were then processed as described below.

### Detection of secreted proteins using Nano-Glo HiBiT Detection System

Secretion cultures were set up and incubated as described above. At the endpoint the OD_600nm_ was measured and volumes containing 3.89×10^9^ bacteria per ml (OD_600nm_ of 1) were centrifuged twice to remove bacteria. The pellets were set aside for further analysis using Nano-Glo HiBiT Lytic Detection System (Promega, N3030). 100 µl of the cleared supernatants were dispensed in a white 96 well plate in triplicate and equilibrated to room temperature. Sterile DM was used as negative control. Nano-Glo HiBiT Extracellular Reagent (Promega, N2420) was prepared following manufactureŕs instructions and 100 µl were dispensed to each sample. The plate was sealed (Thermo Scientific, AB-0580) and the bioluminescence was read in a Synergy H4 microplate reader (Agilent Bio Tek) after 10 min of agitation at room temperature. The reconstitution of the HiBiT and LgBiT fragment in the pellet was analyzed using the Nano-Glo HiBiT Lytic Detection System (Promega, N3030), to show that there is general measurable bioluminescence. For this the beforementioned pellets were resuspended in PBS and centrifuged to remove residual supernatant. The washed pellets were resuspended in 1 ml PBS and 100 µl was dispensed in triplicate into a white 96 well plate. 100 µl PBS were used as negative control. 50 µl of lysis buffer were added to the bacterial suspension and the samples were incubated at room temperature for 10 min to lyse the bacteria. The Nano-Glo HiBiT Lytic reagent was prepared following the manufacturers instructions. 50 µl of reagent were added to each well. The plate was sealed (Thermo Scientific, AB-0580) and the bioluminescence was immediately read in a Synergy H4 microplate reader (Agilent Bio Tek).

### Fractionation of secretion cultures

Secretion cultures were set up and incubated as described before. After incubation a sample equivalent to OD_600nm_ of 0.6 was pelleted and put aside for western blotting. The rest of the cultures were cleared by centrifugation for 20 min at 3000 xg to pellet bacteria. The cleared supernatants were sterile filtered using Nalgene Rapid-Flow filter units with a PES membrane with 0.22 µm pore size (Nalgene, 564-0020). The filtered supernatant was tested for sterility by plating onto LB agar plates. The filtrate was then either subjected to further fractionation to concentrate outer membrane vesicles or subjected directly to TCA-precipitation.

### TCA-precipitation of sterile-filtered supernatant

One volume of 100% trichloroacetic acid (TCA) was added to four volumes of sterile-filtered supernatant and incubated for 1 h at 4°C. The precipitate was sedimented by centrifugation at 10.000 xg for 1 h at 4°C. The supernatant was carefully removed and discarded. The precipitate was rinsed with 0.2 ml acetone. The acetone was removed after centrifugation at 10.000 x g for 1 h at 4°C. 5 µl TRIS-HCl (pH 8.0) was added to equilibrate the pH. The precipitated supernatants were solubilized in appropriate volumes of 2x Laemmli buffer to normalize the samples to ∼1.67×10^9^ CFU in 20 µl based on the OD_600nm_ of the original culture.

### Western blots of fractionated secretion cultures

Bacteria at a density of OD_600nm_ of 0.6 (∼1×10^9^ CFU) were pelleted, washed, and resuspended in 70 μl PBS before heat inactivation for 10 min at 95°C. Standard SDS-sample buffer (Laemmli) was added, and samples were boiled a second time.

Equal volumes were subjected to standard SDS-gel electrophoresis (BioRad, Mini-PROTEAN TGX Precast Protein Gels 4561094) followed by semi-dry western blot. Nitrocellulose membranes (Merck, GE10600001) were blocked with 5% skim milk in 0.1% Tween in PBS (PBS/T) for 1 h at room temperature (RT). Blocked membranes were incubated overnight at 4°C with primary antibody against FLAG-tag (Sigma, F1804) or dsRed (Takara, Living colors dsRED poly Ab, 632496) in blocking buffer. Following 3 wash steps with PBS/T, the membranes were incubated with secondary antibody (CellSignaling, 7076S or 7074S) in blocking buffer for 1 h at RT. Before developing the membranes, they were again washed 3 times with PBS/T. Blots were developed using LumiGlo chemiluminescent substrate (Seracare, 5430) and imaged using the Fusion FX device (Vilber).

### Fractionation of supernatant into soluble and non-soluble fraction using ultracentrifugation

Filtered supernatants were further concentrated by ultracentrifugation to separate soluble and insoluble fraction if needed. After filtration the supernatants were subjected to ultracentrifugation in a Hitachi CS150FNX ultracentrifuge at 100.000 x g for 2 h at 4°C. The supernatant was then carefully removed and subjected to TCA-precipitation as described above, while the OMV-fraction was resuspended in 2x Laemmli normalized to initial ODs of secretion cultures (∼2×10^8^ CFU/μl) to ensure equal loading for western blotting. Samples were boiled for 10 min at 95 °C and subjected to western blotting as described above.

### Prediction of putative signal peptides

Putative signal peptides and N-proximal transmembrane domains (within N-proximal 50 aa) were predicted using SingalP6.0 (https://services.healthtech.dtu.dk/services/SignalP-6.0/) (Almagro Armenteros *et al*. 2019), Phobius (https://phobius.sbc.su.se/) (Käll, Krogh and Sonnhammer 2004), and DeepTMHMM (https://services.healthtech.dtu.dk/services/DeepTMHMM-1.0/) (Hallgren *et al*. 2022).

### Harvesting OMVs for LC-MS/MS proteomics

*Brucella abortus* 544 used for the preparation of OMVs for proteomics was grown as described in (Lannoy *et al*. 2025). In short: six 800 ml cultures were grown for 48 h to stationary phase and inactivated with 0.5 % phenol. Inactivated cultures were centrifuged at 8200 x g at 4°C for 20 min. The pellet was set aside. The supernatant was further concentrated using a Pellicon tangential flow filtration system with a 10 kDa cutoff. Concentrated supernatants were centrifuged at 8200 x g at 4°C for 20 min. The supernatant was frozen at −20°C to aggregate the OMVs. After defrosting the supernatant was ultracentrifuged (47000 x g, 4°C, 3 h) and the pellet containing the OMVs was resuspended in deionized water and dialysed at 4°C with deionized water for 3 days. Samples were stored at −80°C and then OMVs and bacterial pellets were lyophilized by freeze drying (Telstar Cryodos 50).

### Preparation of OMVs for LC-MS/MS proteomics

Freeze dryed OMVs were resuspended in ultrapure water (0.4 % (w/v)) and treated with Triton ×100 (1% (w/v)) for 1 h at 37°C. The suspension was centrifuged for 3 min at 5.000 x g. 10 µl of 100% TCA were added to 900 µl supernatant, vortexed, and incubated for 30 min on ice. Lysates were then centrifuged at 10.000 xg for 15 min at 4°C and washed with ice-cold acetone. The resulting protein pellet was resuspended in 100 µl 2% SDS in ddH_2_O and heated for 5 min at 95 °C. The protein concentrations were measured by the Lowry test (Lowry *et al*. 1951) prior to protein digestion.

### Protein digestion for LC-MS/MS proteomics

Protein Samples were digested using the Filter Aided Sample Preparation protocol (Wiśniewski *et al*. 2009). Briefly, 30 KDa molecular weight cut-off filters (Millipore, Amicon Ultra), were washed using 100 µl 0.1% Formic Acid (FA) and centrifuged at 14500 rpm for 15 min. Next 40 µg of protein lysate in urea buffer (8M urea in 0.1M Tris at pH 8.5) was added to the filter and centrifuged at 14500 rpm for 15 min. The filter was washed 3 times using 200 µl urea buffer, discarding the filtrate between each centrifugation step. Proteins were reduced by adding 100 µl 8 mM DTT to each filter then mixed for 1 min at 400 rpm with a thermomixer, and then incubated at 24 for 15 minutes before centrifugation at 14500 rpm for 15 min. After discarding the filtrate, the filter was washed with 100 µl urea buffer. Proteins were then alkylated by adding 100 µl 50 mM iodoacetamide (IAA), in urea buffer, and mixing at 400 rpm on a thermomixer before incubating in the dark for 20 min at 24°C. Filters were then centrifuged at 14500 rpm for 15 min, and washed with 100 µl urea buffer.

Samples were then washed 3 times with 100 µl of 50 mM ammonium bicarbonate buffer (ABC, in ultrapure water). Following this, 0.8 µg of MS grade trypsin (in 80 µl ABC buffer) was added, before overnight incubation at 24°C in a high humidity environment.

Following overnight digestion, peptides were eluted by centrifugation at 14’500 rpm for 10 min, into clean LoBind centrifuge tubes, with an additional 40 µl ABC buffer added to the filter. Penultimately, 10% Trifluoroacetic acid (TFA) was added to a final concentration of 0.2% TFA. Samples were subsequently dried in a SpeedVac, and up to 20 µl were transferred to an injection vial.

### Proteomics data acquisition

The digest was analyzed using nano-LC-ESI-MS/MS tims TOF Pro (Bruker, Billerica, MA, USA) coupled with an UHPLC nanoElute2 (Bruker).

Peptides were separated by nanoUHPLC (nanoElute2, Bruker) on a 75 μm ID, 25 cm C18 column with integrated CaptiveSpray insert (Aurora, ionopticks, Melbourne) at a flow rate of 200 nl/min, at 50°C. LC mobile phase A (0.1% FA in water) and B (Acetonitrile with 0.1% FA). Samples were loaded directly on the analytical column at a constant pressure of 800 bar. The 1 µl of digest was injected, and the organic content of the mobile phase was increased linearly from 2% B to 15% in 22 min, from 15% B to 35% in 38 min, from 35% B to 85% in 3 min. Data acquisition on the tims TOF Pro was performed using Hystar 6.1 and time Control 2.0. tims TOF Pro data were acquired using 160 ms TIMS accumulation time, mobility (1/K0) range from 0.75 to 1.42 Vs/cm². Mass-spectrometric analysis were carried out using the parallel accumulation serial fragmentation (PASEF) (Meier *et al*. 2018) acquisition method. One MS spectra followed by six PASEF MSMS spectra per total cycle of 1.16 s. The mass spectrometry proteomics data have been deposited to the ProteomeXchange Consortium via the PRIDE (Perez-Riverol *et al*. 2023) partner repository with the dataset identifier PXD067564.

### Proteomics Database Search

Resulting .d files were searched with FragPipe using msFragger version 4.1 and IonQuant version 1.10.27, with the default parameters for the LFQ-MBR workflow (Yu, Haynes and Nesvizhskii 2021; Yu, Deng and Nesvizhskii 2025). Briefly, precursor and fragment mass tolerance were set to 20 ppm. The *in silico* peptide digestion parameters were set to use strict trypsin rules, with a maximum of 2 missed cleavages. Peptides between 7 and 50 amino acids long, and masses between 500 and 5000 daltons, peptides were generated. Methionine oxidation and N-terminal acetylation were set to variable modifications and cystein carbomidomethylation was set as a fixed modification. All results were filtered to 0.01% FDR at the precursor and protein-level. All files were searched against the UniProt (The UniProt Consortium *et al*. 2023) fasta database for *B. abortus (strain 2308) -* UP000002719 (downloaded on 21 July 2025). Proteins were considered “identified” if proteotypic peptides for a given protein below the FDR cut-off were identified in at least one sample. Proteins were considered “quantified” if for a given protein in at least 2 samples the MaxLFQ Intensity was greater than 0. i.e, there was at least 1 shared peptide, between samples.

### Bioinformatic analyses

#### OMV vs whole cell rank difference

Quantified proteins in both the OMV and whole cell fractions were ranked from highest to lowest according to their MaxLFQ intensities in each fraction. The rank difference was calculated by subtracting the OMV rank from the whole cell rank. A positive rank difference indicates relative enrichment in OMVs and a negative rank difference indicates a relative enrichment in whole cells. We also calculated the log2 Fold Change between proteins quantified in both OMVs and Whole-cells.

#### Orthogroup mapping

Orthogroups were generated using OrthoFinder version 2.5.5 (Emms and Kelly 2019), with default settings, to match *Brucella* proteins between species. Which allowed us to directly compare published proteomes at the orthogroup level. The protein sequence databases for *Brucella melitensis* biotype 1 (UP000000419), *Brucella suis* biovar 1 (strain 1330) (UP000007104) and *Brucella abortus (strain 2308)* (UP000002719), were downloaded from uniprot on 21 July 2025. The orthogroup mappings can be found in supplementary Data S1.

#### Statistical analysis

Graphs were created with GraphPad Prism 8. Statistical analysis was performed using GraphPad Prism with ordinary one-way ANOVA. The number of independent replicates is indicated in the figure legends as n.

## Results

### Analysis of *Brucella abortus* whole-cell and OMV proteomes

Given that various *Brucella* species produce OMVs (Avila-Calderón et al., 2012, 2020; Pollak et al., 2012; Araiza-Villanueva et al., 2019; Socorro Ruiz-Palma et al., 2021) and that these vesicles have been shown to modulate immune responses, induce cytoskeleton rearrangements, and promote bacterial internalization *in cellulo* (Pollak et al., 2012; Avila-Calderón et al., 2020), we sought to determine if *Brucella* effector proteins localize to OMVs, which would suggest that OMVs represent a previously uncharacterized route for effector secretion.

To investigate the protein content and the relative abundance of described effector proteins in OMVs, we purified OMVs from *B. abortus* as described in (Lannoy *et al*. 2025). The OMV- and whole-cell-fractions were analyzed by LC-MS/MS. In total, we identified 18’016 distinct *B. abortus* peptide sequences corresponding to 1’947 proteins at an FDR threshold of 0.01%. Among these, 1’186 proteins were identified in both OMV and whole cell-fractions, and 1’153 quantified in both. Here, we distinguish between identification and quantification based on the number of unique peptides detected: proteins were considered quantified only if at least one shared peptide was identified across two of three samples. 25 proteins were identified exclusively in the OMV-fractions, of which 21 were quantified, while 717 proteins were identified only in the whole cell-fraction, with 372 quantified. The large number of proteins detected exclusively in the whole cell-fraction, but absent from the OMV-fraction, argues against widespread bacterial lysis as the source of OMV-associated proteins.

Tables of all identifications and quantifications, as well as associated scores are available in the supplement (Data S2 and S3), including tables of exclusively identified and quantified proteins.

### Inner membrane and cytosolic proteins are depleted from the OMV fraction

To assess the quality of our OMV-preparation, we ranked all proteins quantified at least once in OMVs and whole-cell samples by abundance and calculated their log_2_ fold changes (Data S4). In this analysis, a positive rank denotes enrichment in the OMV-fraction, whereas a negative rank denotes higher abundance in whole-cells. As reference groups, we then selected 19 cytosolic, 10 inner membrane-associated, 10 outer membrane-associated, and 11 periplasmic proteins to evaluate sample purity. Inner membrane proteins were strongly depleted from the OMV-containing fractions (9 of 10 proteins with negative rank differences and all with negative log_2_ fold changes) (Fig. 1A). Cytosolic proteins were also mostly depleted from the OMV-containing fraction, with most displaying negative rank differences and a log_2_ fold changes (Fig. 1A). By contrast, outer membrane and periplasmic proteins were more evenly distributed, with 6 of 10 outer membrane and 6 of 11 periplasmic proteins showing depletion (Fig. 1B). Among the outer membrane proteins, the adhesin BmaC (Bialer *et al*. 2021) was highly enriched in the OMV-containing fraction (log_2_ fold change = 2.56), whereas Omp2b and Omp28 were strongly depleted (log_2_ fold change of −1.91 and −1.84, respectively) (Fig. 1B and Data S4), which is consistent with Omp2b being covalentaly linked to peptidoglycan and might be true for the other depleted Omps (Godessart *et al*. 2021).

**Figure 1:**
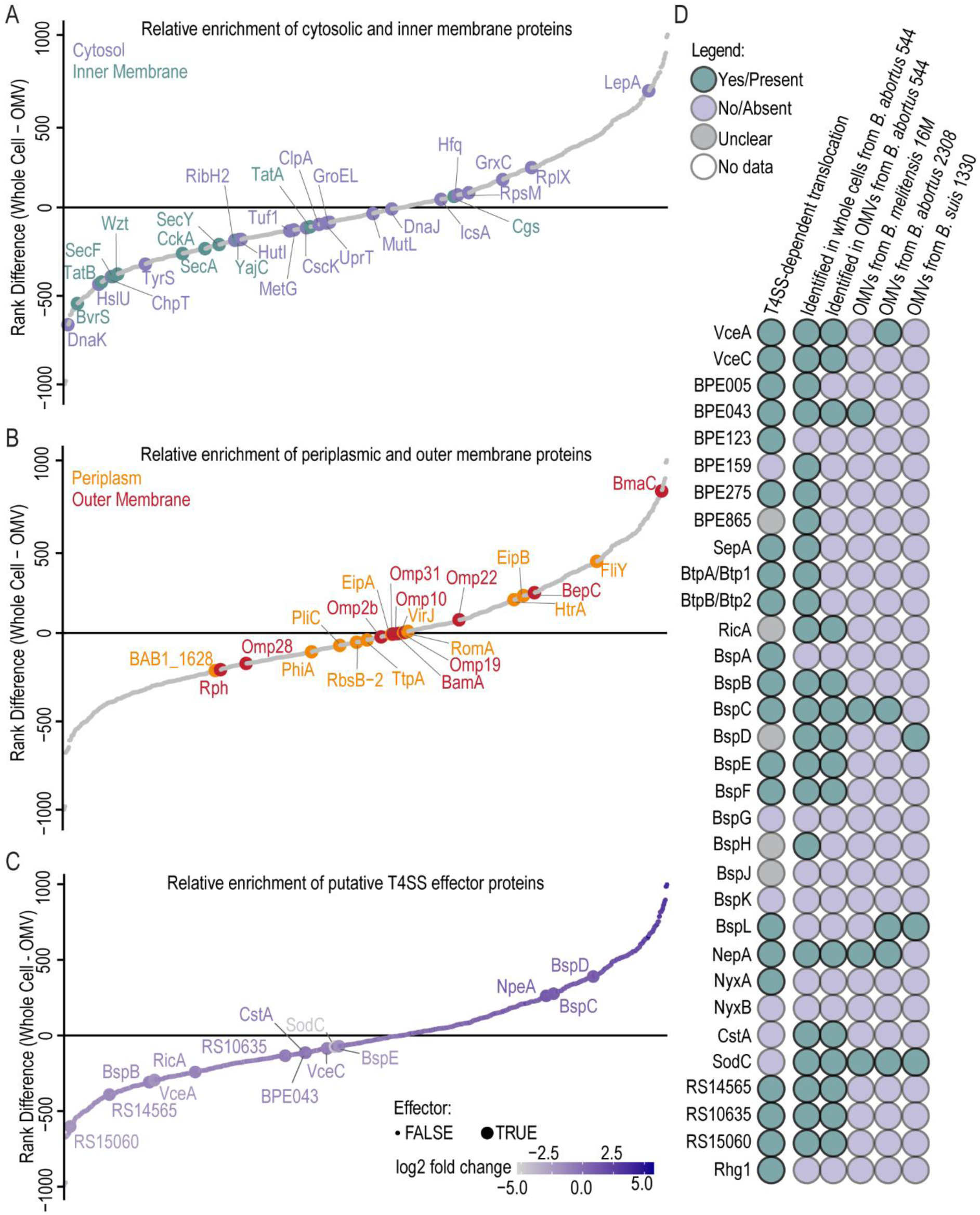
A subset of putative *Brucella* effectors associates with OMVs. **(A,B)** Proteins quantified at least once in OMV and whole-cell fractions of *B. abortus* 544 were ranked by abundance in descending order. The difference in rank (whole-cell rank - OMV rank) was plotted, with positive values indicating enrichment in OMVs and negative values enrichment in the whole-cells. To assess OMV preparation quality, the relative abundance of 19 cytosolic, 10 inner membrane-associated, 11 periplasmic proteins, and 10 outer membrane proteins was evaluated. **(C)** Effectors quantified at least once in OMV and whole-cell fractions were ranked by abundance in descending order. **(D)** In *B. abortus* 544, 23 described effector proteins were identified in the whole-cell fraction and 15 in the OMV-fraction. Comparison with published datasets revealed that a subset of putative *Brucella* effector proteins associates with OMVs in different *Brucella* species. OMV proteomes of *B. melitensis* 16M (Avila-Calderón *et al*. 2020), *B. abortus* 2308 (Araiza-Villanueva *et al*. 2019), and *B. suis* 1330 (Socorro Ruiz-Palma *et al*. 2021) were analyzed. T4SS-dependency was inferred from the literature.

These results indicate minor cytosolic contamination, which is expected as bacteria were grown to stationary phase and some lysis may have occured, or alternatively that certain cytoplasmic proteins selectively associate with OMVs by an unknown mechanism. Nevertheless, the strong depletion of inner membrane proteins from the OMV-preparations, together with the exclusive identification of 717 proteins in whole-cell fractions (see above), supports that the OMV preparation was relatively clean.

### A subset of putative *Brucella* effectors localize to OMVs

Next, we investigated whether described effector proteins localize to OMVs and whether they are enriched in these fractions. Of the 32 described effectors, 23 were identified in the whole-cell fraction, and 15 of these were also detected in the OMV-containing fractions, indicating that a subset of effectors associates with OMVs (Fig. 1C and D). BspF was detected only once in both fractions and was therefore not quantified. Similary, BPE275 and BtpB were detected only once in the whole-cell fraction and were excluded from quantification.

To assess enrichment, proteins quantified at least once in OMV and whole-cell factions were ranked by abundance (Fig. 1C, Data S4 and S5). Log2 fold-change analysis revealed a strong correlation with rank differences (Fig. S1, Data S4). In this analysis, NpeA, BspC, and BspD were enriched in OMVs releative to whole-cells, with a rank difference of 262, 276 and 390, respectively. We further compared the protein contents of *B. abortus* OMVs from this study with previously published datasets of OMVs from *B. abortus*, *B. melitensis*, and *B. suis* (Avila-Calderón *et al*. 2012; Araiza-Villanueva *et al*. 2019; Socorro Ruiz-Palma *et al*. 2021) (Fig. 1D, Data S6). Across at least two of the four datasets analyzed, VceA, BspC, BspD, BspL, BPE043, NpeA, and SodC were consistently associated with OMVs (Data S6). Of these, five proteins (VceA, BPE043, BspC, BspL, NpeA) have been reported as T4SS-dependent effectors, one (BspD) has an undefined translocation mechanism, and one (SodC) is T4SS-independent (De Jong *et al*. 2008; Marchesini *et al*. 2011; Myeni *et al*. 2013; Liu *et al*. 2018; Luizet *et al*. 2021; Giménez *et al*. 2024). Notably, the *Brucella* species have been cultured under different conditions and OMVs were enriched using different methods, suggesting that effector association with OMVs is robust and not an experimental artifact.

Together, these results demonstrate that effector proteins of *Brucella* can localize to OMVs and suggest that OMVs may served as an alternative pathway for the secretion of virulence factors, a phenomenon likely overlooked in the past due to methodological limitations.

### A subset of putative effector proteins encode predicted signal peptides and transmembrane domains

Next we asked whether OMV-associated effector proteins share any structural features such as signal peptides (SPs) or transmembrane domains (TMs). Canonical SPs consist of a positively charged N-terminal region, a central hydrophobic h-region, and a polar C-terminal region containing the cleavage site for signal peptidase processing (von Heijne 1990). Proteins with classical SPs are typically secreted to the periplasm via the conserved SecYEG translocon of the inner membrane, with assistance from SRP/YidC or SecA (Walter, Ibrahimi and Blobel 1981; Gilmore, Blobel and Walter 1982).

*In silico* analysis with SignalP6.0 (Almagro Armenteros *et al*. 2019), Phobius (Käll, Krogh and Sonnhammer 2004) and DeepTMHMM (Hallgren *et al*. 2022) predicted SPs and N-proximal TMs in a subset of *Brucella* effector proteins (Fig. 2A, Table S4 and S5). VceA, BspC, BspL, and SodC displayed strong SP predictions, while NpeA carried a predicted lipoprotein SP, as previously reported (Giménez *et al*. 2024). For BPE123, RS10635, and BspB, Phobius predicted SPs that were not revealed by SignalP6.0, whereas DeepTMHMM instead predicted N-terminal TMs for these proteins. N-proximal TMs were consistently predicted for VceC, BPE159, BPE275, BspA, BspD, BspK, and Rhg1. In total, 15 of 32 putative secreted or translocated *Brucella* effector proteins encoded predicted N-terminal features. Of particular note, among the seven effector proteins localized to OMVs in at least two independent datasets (Fig. 1D), five (VceA, BspC, BspL, NpeA, SodC) carried predicted SPs, one (BspD) carried a predicted TM, and one (BPE043) lacked any predicted N-terminal feature (Fig. 2A). Importantly, all effector proteins with predicted SPs were found to localize to OMVs but periplasmic proteins were not generally enriched in the OMVs, strongly suggesting that *Brucella* employs OMVs as vehicles for SP-dependent secretion of virulence factors.

**Figure 2:**
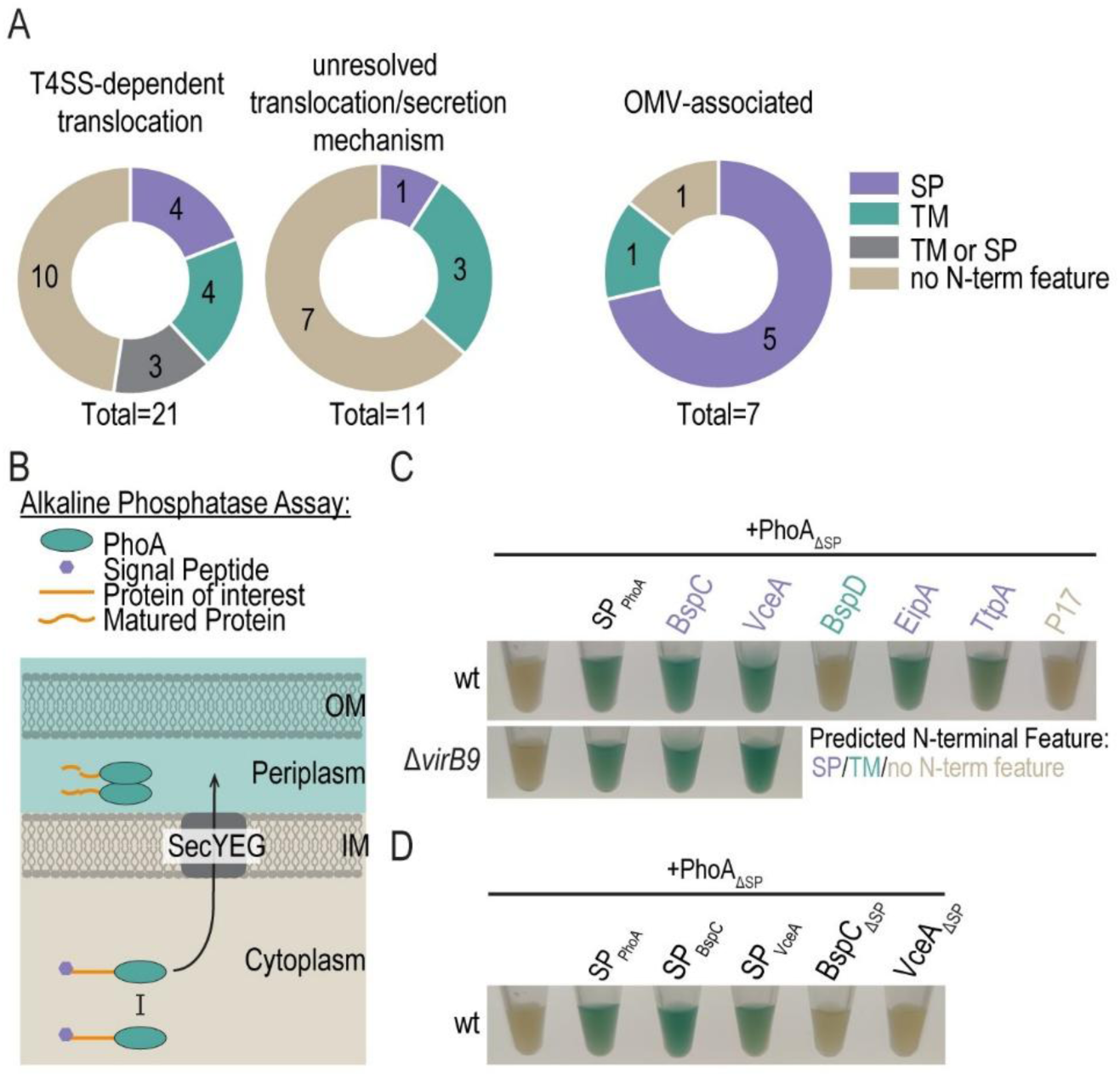
The OMV-associated effectors VceA and BspC encode functional signal peptides. **(A)** *In silico* prediction of N-terminal features of secreted and translocated *Brucella* proteins identified three groups: proteins with predicted N-proximal transmembrane domains (TM), proteins with signal peptides (SP), and proteins with ambiguous predictions (TM or SP or no N-terminal feature). **(B)** Schematic representation of the alkaline phosphatase (PhoA) assay. PhoA dimerizes and becomes active only in non-reducing environments (periplasm or extracellular milieu), but remains inactive in the reducing cytoplasm. Signal peptides are cleaved during export via the SecYEG translocon. OM = outer membrane, IM = inner membrane, SecYEG = Sec translocon. **(C,D)** *B. microti* strains expressing the indicated PhoA-fusion proteins were grown to exponential phase in TSB. Cells corresponding to an OD_600nm_ of 2 (per 1 ml culture) were pelleted, resuspended in 100 µl fresh TSB supplemented with 200 µg/ml BCIP, and incubated for 1-2 h at 37°C before imaging. Predicted N-terminal features are indicated by colours. Shown is one representative experiment of two independent replicates.

### The putative effector proteins BspC and VceA are exported from the cytoplasm to the periplasm via the SecYEG-translocon

We further investigated the functionality of the predicted SPs. For this purpose, we selected VceA, BspC, and BspD for further validation because (i) the SP-predictions for BspC and VceA were particularly strong, while BspD has a predicted TM, and (ii) they represent different classes of putative effectors: those which can localize to OMVs (VceA), and those which are enriched in OMVs (BspC, BspD).

We used the *Escherichia coli* protein alkaline phosphatase (PhoA) as a reporter (Kumamoto, Oliver and Beckwith 1984; Kim and Harold 1991; Herrou *et al*. 2019a, 2019b) in combination with *Brucella microti*, a Biosafety-level 2-compatible surrogate for human-pathogenic *Brucella* species. PhoA becomes active only when exported to the non-reducing environment of the periplasm via the SecYEG-translocon, where it catalyzes the hydrolysis of organic phosphate esters (Kumamoto, Oliver and Beckwith 1984; Kim and Harold 1991). This activity can be visualized colorimetrically using substrates such as 5-Bromo-4-chloro-3-indolyl phosphate disodium salt (BCIP), which is colorless but turns blue upon phosphate hydrolysis and poorly penetrates the inner membrane (Fig. 2B).

PhoA-fusions were expressed from a plasmid under an aTc-inducible promoter in *Brucella*. Replacement of the native PhoA SP (PhoA^M1-A21^, hereafter SP_PhoA_) with a Flag-tag (yielding Flag-PhoA^R22-K471^, hereafter PhoA_ΔSP_), abolished periplasmic export while maintaining expression (Fig. 2C, Fig. S2). Export was reconstituted by fusing SP_PhoA_ to the N-terminus of PhoA_ΔSP_ (Fig. 2C). Control fusions confirmed the localization: the cytoplasmic protein P17 showed no BCIP conversion, whereas the periplasmic proteins EipA and TtpA resulted in blue coloration consistent with periplasmic localization (Fig. 2C). Using this assay, we found that the putative T4SS effectors VceA and BspC localized to the periplasm in a SP-dependent, and thus Sec-dependent manner (Fig. 2C and D). In contrast, the envelope integrity protein and putative effector BspD (Myeni *et al*. 2013; Ketterer *et al*. 2024) was not exported, consistent with the absence of a predicted SP and presence of a predicted N-terminal TM (Table S5, Fig. 2C). We further tested whether export of VceA and BspC to the periplasm was T4SS-independent, as the association to OMVs suggests, and indeed, export was unaffected in a *B. microti* Δ*virB9* mutant, which cannot assemble a functional T4SS (O’Callaghan *et al*. 1999).

In summary, VceA and BspC are exported from the cytoplasm to the periplasm in a SP-dependent, T4SS-independent manner via the SecYEG translocon.

### VceA and BspC are secreted to the culture supernatant in a SP-dependent and T4SS-independent manner

We next tested whether the T4SS effectors VceA and BspC (De Jong *et al*. 2008; Myeni *et al*. 2013), which are exported to the periplasm via the SecYEG-translocon (Fig. 2C) and are associated to OMVs (Fig. 1C and D), could be secreted into the culture supernatant in a SP-dependent manner.

For this purpose, we employed a split NanoGlo (NGlo) luciferase assay, previously applied to monitor effector protein translocation in host cells by the T3SS of *Salmonella* and T4SS of *Bartonella* (Westerhausen *et al*. 2020; Fromm *et al*. 2022, 2024), as well as to quantify OMV production in culture supernatants of *Bacteroides* (Pardue *et al*. 2024). Proteins of interest were fused to the HiBiT fragment and expressed in *B. abortus*, while LgBiT was supplemented to cleared supernatants (to detect secreted proteins) and to lysed bacterial pellets (to measure input). *B. abortus* cultures were grown to an OD_600nm_ of 1.0-1.4 in a defined medium (DM) as described previously (De Barsy *et al*. 2011) and pelleted, and supernatants were clarified by additional centrifugation.

As expected, the cytoplasmic protein P17, which was strongly depleted from our OMV-preparation (rank difference: −184, log2 fold change: −1,43) (Data S4), was not detected in supernatants, confirming the absence of bacterial lysis (Fig. 3A and Fig. S3A). Similarly, the effector protein BPE123, not associated to OMVs (Fig. 1B), was undectectable in the culture supernatant. The periplasmic protein EipA (Herrou *et al*. 2019a), which was depleted from our OMV-preparation (rank difference of −7, log2 fold change of −0.5) (Data S4), also did not appear in supernatants (Fig. 3A and Fig. S3A), showing that overexpression of periplasmic proteins alone does not drive OMV release. By contrast, both VceA and BspC were secreted in a SP-dependent manner (Fig. 3A Fig. S3A), with VceA showing a ∼5-6 fold enrichment in the supernatant relative to the pellet. Secretion of BspC and VceA was also detected in *B. microti* (Fig. 3C and Fig. S3B), indicating that this phenomenon is conserved across *Brucella* species. However, while VceA was highly enriched in the OMV fraction from the *B. abortus* 2308 culture it was not enriched in the OMV fraction of the *B. microti* culture relative to pellet (Fig. 3A and C), indicating potential differences between species and strains in respect to sorting and packaging, as we did not observe enrichment of VceA in the OMV fraction from *B. abortus* 544 cultures. Notably, *B. abortus* 544 was grown to stationary phase in rich medium, while *B. abortus* 2308 and *B. microti* were grown to exponential phase in defined medium, which might also account for some of the observed differences.

**Figure 3:**
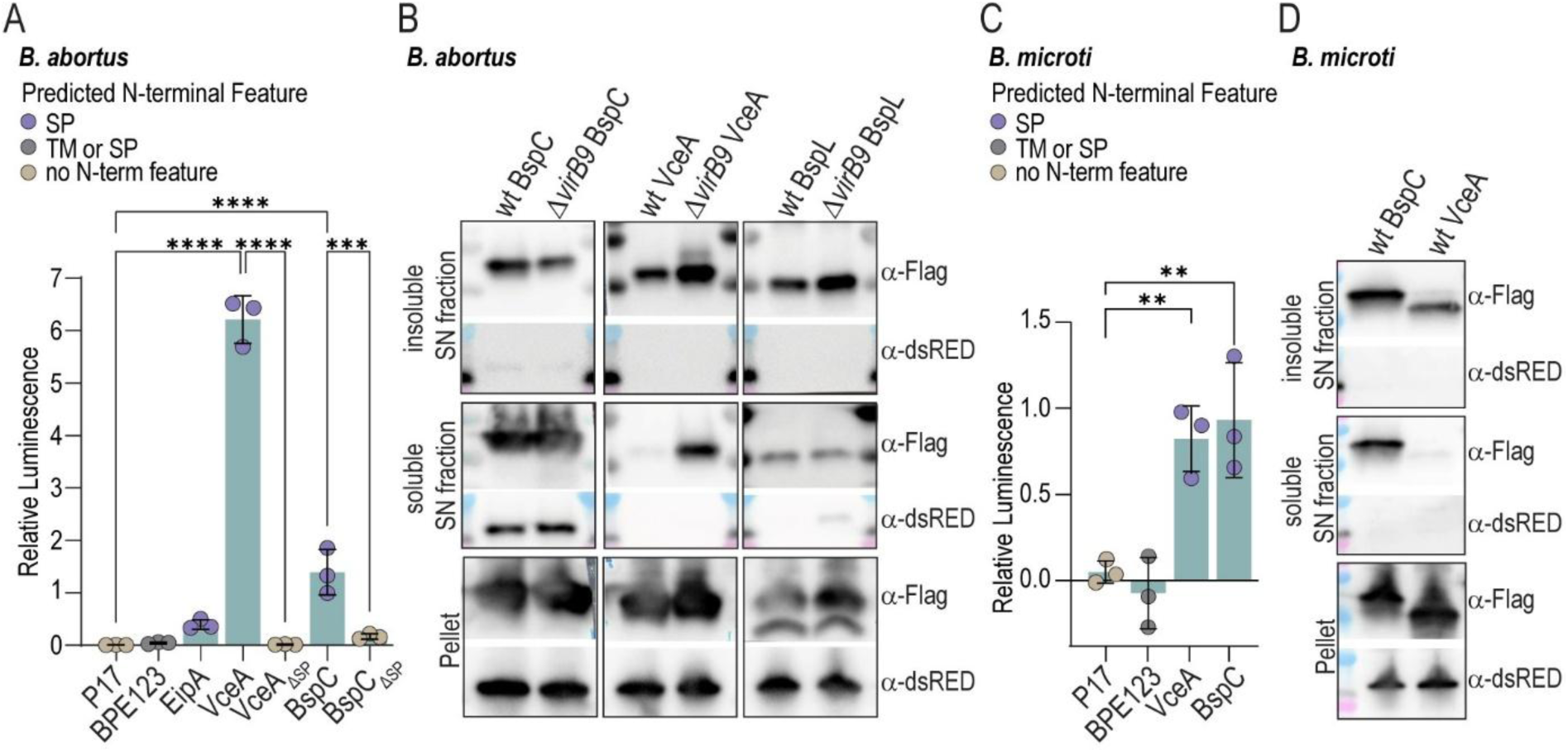
*Brucella spp.* secrete VceA, BspC, and BspL to the culture supernatant. **(A)** *B. abortus* strains expressing the indicated HiBiT fusion proteins were grown to exponential phase in 2YT. Secretion cultures were incubated for 17 h in defined medium (DM), after which bacteria were pelleted and cleared supernatants collected. Bacterial pellets were lysed, and both fractions were supplemented with LgBiT and substrate. Bioluminescence was measured using a plate reader, and relative luminescence was calculated as the ratio of cleared supernatant versus lysed pellet signals. Ordinary one-way ANOVA; n=3. Predicted N-terminal features are indicated by colours. **(B)** Axenic secretion assay with *B. abortus* evaluating localization of BspC, VceA, and BspL in soluble (OMV-depleted) versus insoluble (OMV-containing) supernatant fractions. Sterile-filtered supernatants were fractionated by ultracentrifugation, and both fractions, alongside bacterial pellets, were analyzed by SDS-PAGE and western blotting using α-Flag and α-dsRED antibodies. Shown is one exemplary blot of 3 independent replicates. **(C)** *B. microti* strains expressing the indicated HiBiT fusion proteins were analyzed as described in (A), however, due to their faster growth rate, the incubation time in DM was reduced to 8h. n=3. **(D)** Axenic secretion assay with *B. microti* for evaluating localization of BspC and VceA as described in (B). Shown is one exemplary blot of 3 independent replicates.

To confirm OMV association, sterile-filtered supernatants were fractionated by ultracentrifugation into soluble (OMV-depleted) and insoluble (OMV-containing) fractions and analyzed by SDS-PAGE and western blotting (Fig. 3B). VceA as well as BspL (Luizet *et al*. 2021) were enriched in the insoluble, OMV-containing fraction without evidence of bacterial lysis, as assessed by absence of cytosolic dsRED, which was co-expressed from the same plasmid. In contrast, BspC overexpression resulted in partial lysis, as indicated by the presence of both dsRED and BspC in the soluble fraction (Fig. 3B).

To assess T4SS dependency, we repeated the assay in a *B. abortus* Δ*virB9* mutant lacking a functional T4SS (Fig. 3B). Secretion of VceA and BspC remained unaffected, confirming T4SS independence. Interestingly, VceA was additionally detected in the soluble fraction of the *ΔvirB9* mutant in the absence of dsRED, though its enrichment in the insoluble fraction was preserved (Fig. 3B). Finally, BspC and VceA were again detected in OMV-containing fractions of *B. microti* (Fig. 3D), confirming that effector localization to OMVs is conserved across *Brucella* species.

Together, these data demonstrate that a subset of described T4SS effector proteins, including VceA and BspC, are secreted into the culture supernatant in a SP-dependent, T4SS-independent manner and associate with OMVs, a process conserved among *Brucella* species.

## Discussion

One of the major challenges *Brucella* faces toward the establishment of its replicative niche in the ER is the manipulation of host cell processes during intracellular trafficking. Although *Brucella* was first described over a century ago by Sir David Bruce, the precise molecular mechanisms underlying its intracellular lifestyle remain incompletely understood. It is well established that *Brucella* depends on its T4SS for intracellular replication, and an increasing number of putative T4SS effectors have been identified in recent years. However, how these effector proteins are translocated into host cells and how they contribute to pathogenesis remains poorly defined. Translocation assay data are often inconsistent (Myeni *et al*. 2013), and the existence of T4SS-independent effector proteins suggests that additional secretion mechanisms are involved.

Several *Brucella* species are known to produce OMVs and OMV-resembling structures can be detected in the vicinity of bacteria during monocyte infection (Boigegrain *et al*. 2004; Avila-Calderón *et al*. 2012, 2020; Pollak *et al*. 2012; Araiza-Villanueva *et al*. 2019; Socorro Ruiz-Palma *et al*. 2021). Further, *B. melitensis* OMVs induced TNFα and IL-6 secretion and cytoskeletal rearrangements in PBMCs, but not IL-17 secretion, apoptosis, DNA damage, or proliferation; these effects were independent of LPS phenotype (Avila-Calderón *et al*. 2020). This demonstrates that *Brucella* OMVs can modulate immune responses, although the underlying factors and *in vivo* relevance remain unclear. We therefore hypothesized that OMVs might represent an alternative route for effector protein secretion. Indeed, our study demonstrates that several known and putative T4SS effectors are consistently associated with outer membrane vesicles produced by *Brucella* spp.. Strinkingly, especially effector proteins with predicted signal peptides were detected in OMVs, suggesting that SPs may be a prerequisite for OMV targeting. However, periplasmic proteins were not generally enriched in OMVs compared to whole-cells, implying the involvement of additional sorting mechanisms.

In our analysis, 15 of 32 described effector proteins were detected in *B. abortus* OMVs grown to stationary phase in rich medium. Comparative proteomics with previously published datasets from *B. melitensis*, *B. abortus*, and *B. suis* revealed seven effectors (VceA, BspC, BspD, BspL, BPE043, NpeA, SodC) consistently present in at least two of four datasets, suggesting robust OMV association. NpeA, BspC, and BspD were enriched in OMVs relative to whole-cells in our dataset. Importantly, these findings were made under axenic growth conditions, and effector expression is likely to differ during host infection, potentially increasing the repertoire of OMV-associated proteins. It will therefore be important to study the influence of infection-mimicking conditions (such as acidic pH and nutrient limitation) on OMV-cargo composition in the future. The strong enrichment of certain proteins in the OMVs compared to the whole-cells and the association of several effector proteins with OMVs of different *Brucella* species, favors a model of controlled vesicle biogenesis rather than passive blebbing and supports the hypothesis that OMVs serve as specific vehicles for virulence factor secretion. To this end it will be important to explore if the OMVs of *Brucella* spp. mediate direct translocation of effectors into host cells by adapting and utilizing reporter assays such as the TEM1 β-lactamase assay. However, since we lack knowledge about OMV-biogenesis in *Brucella* as well as an understanding at which infection stages OMVs might be utilized devising such an assay is not trivial. *Brucella,* as a facultative intracellular pathogen, may employ OMVs at defined infection stages. OMVs could be produced within host membrane-bound compartment as described for *Salmonella enterica* serovar Typhimurium, and released to the host cell cytosol (Yoon *et al*. 2011; Guidi *et al*. 2013). Alternatively, OMV release could occur upon T4SS-dependent damage of the *Brucella*-containing vacuole (BCV), observed as early as 6 h post-infection (Costa Franco *et al*. 2018; Tana *et al*. 2021; Hiyoshi *et al*. 2022). Such BCV damage could release luminal components, including OMVs, or even bacteria into the cytoplasm. Whether *Brucella* produces OMVs *in cellulo* or *in vivo* remains an open question.

For other pathogens, the role of OMVs in virulence is well documented. *Helicobacter pylori* OMVs deliver VacA toxin, detectable in gastric biopsies (Fiocca *et al*. 1999). *S. enterica* serovar Typhimurium produces toxin-containing OMVs inside host cells, which are secreted and taken up by neighboring cells (Guidi *et al*. 2013). *Coxiella burnetii* appears to release periplasmic proteins into host cell cytosol, and OMVs have been proposed as transport vehicles (Stead *et al*. 2013). *Legionella pneumophila* OMVs inhibit phagosome-lysosome fusion (Fernandez-Moreira, Helbig and Swanson 2006) and promote replication when host cells are pretreated with OMVs (Jung *et al*. 2016). Interestingly, *Legionella* T4SS mutants remain viable but do not follow normal intracellular trafficking, raising the possibility that OMVs partially compensate for T4SS function (Fernandez-Moreira, Helbig and Swanson 2006). Similarly, *Brucella* T4SS mutants remain viable but fail to replicate or follow canonical intracellular trafficking (Comerci *et al*. 2001; Celli *et al*. 2003). This raises the possibility that additional secretion mechanisms, including OMV production inside host cells, contribute to intracellular survival.

## Acknowledgments

Calculations were performed at sciCORE (http://scicore.unibas.ch/) scientific computing center at University of Basel. This work was supported by the Swiss National Science Foundation (SNSF, www.snf.ch) grant 320030-231940 (10003225) to C.D., by PDR grants T.0058.20 and T.0068.24 from FRS-FNRS, as well as Concerted Research Action 17/22-087 and 22/27-128 from the Fédération Wallonie-Bruxelles to X.D.B., and by a FRIA (FNRS) PhD fellowship to A.L..

## Author contributions

M.K., C.D.: Conceptualization, Supervision, Project administration. M.K., N.G.: Visualization, Data curation, Methodology, Formal analysis, Validation. M.K., P.C., A.V., J.B.: Investigation. N.G.: Software. A.L., M.D.: Resources. M.K., N.G., C.D.: Writing – Original Draft. M.K., C.D., M.Q., X.D.B.: Writing – Review & Editing. C.D., X.D.B., A.L.: Funding acquisition.

