## SupplementaryInformation for "A subset of type 4 secretion system effectors of *Brucella* spp. associates to outer membrane vesicles"

### Supplementary Information

Table S1: Strains used in this study.

| Strain | Relevant characteristics | Reference/Source | Description |
|---|---|---|---|
| ***Escherichia coli*** | | | |
| Stellar | *F-, endA1, supE44, thi-1, recA1, relA1, gyrA96, phoA, Φ80d lacZΔ M15, Δ(lacZYA-argF) U169, Δ(mrr-hsdRMS-mcrBC), ΔmcrA, λ-* | Takara bio | Standard cloning strain |
| JKE201 | MFDpir Δ*mcrA* Δ*(mrr-hsdRMS-mcrBC) aac(3)IV::lacIq* | (Harms *et al.* 2017) | Donor strain for conjugation |
| ***Brucella abortus*** | | | |
| | *Brucella abortus 544* | J-M. Verger, INRA, Tours | |
| RCBr01 | *Brucella abortus* 2308 | Kind gift of J. P. Gorvel, Centre d'Immunologie de Marseille-Luminy (CIML) (FR) | |
| RCBr03 | *Brucella abortus* 2308 Δ*virB9* | Kind gift of J. Celli, Washington State University (USA) (Celli, Salcedo and Gorvel 2005) | |
| MKBr144 | *B. abortus* p*bspC-hibit* | This study | |
| MKBr146 | *B. abortus* Δ*virB9* p*bspC-hibit* | This study | |
| MKBr150 | *B. abortus* p*ha.bspC*$_{\Delta SP}$*-hibit* | This study | |
| MKBr156 | *B. abortus* p*bpe123-hibit* | This study | |
| MKBr160 | *B. abortus* p*vceA-hibit* | This study | |
| MKBr162 | *B. abortus* Δ*virB9* p*vceA-hibit* | This study | |
| MKBr164 | *B. abortus* p*vceA*$_{\Delta SP}$*-hibit* | This study | |
| MKBr138 | *B. abortus* p*p17-hibit* | This study | |
| MKBr152 | *B. abortus* p*eipA-hibit* | This study | |
| MKBr166 | *B. abortus* p*bspL-hibit* | This study | |
| MKBr168 | *B. abortus* Δ*virB9* p*bspL-hibit* | This study | |
| ***Brucella microti*** | | | |
| MKBMi103 | *Brucella microti* CCM 4915T | Kind gift of Sascha Al Dahouk, Product Hygiene and Disinfection Strategies at Bundesinstitut für Risikobewertung Berlin (GER) | |
| MKBMi139 | *B. microti* CCM 4915T Δ*virB9* | This study | |
| MKBMi128 | *B. microti* p*phoA*$_{\Delta SP}$ | This study | |
| MKBMi251 | *B. microti* Δ*virB9* p*phoA*$_{\Delta SP}$ | This study | |
| MKBMi130 | *B. microti* p*SP*$_{phoA}$-*phoA*$_{\Delta SP}$ | This study | |
| MKBMi252 | *B. microti* Δ*virB9* p*SP*$_{phoA}$-*phoA*$_{\Delta SP}$ | This study | |
| MKBMi134 | *B. microti* p*bspD-phoA*$_{\Delta SP}$ | This study | |
| MKBMi144 | *B. microti* p*bspC-phoA*$_{\Delta SP}$ | This study | |
| MKBMi253 | *B. microti* Δ*virB9* p*bspC-phoA*$_{\Delta SP}$ | This study | |
| MKBMi145 | *B. microti* p*SP*$_{bspC}$-*phoA*$_{\Delta SP}$ | This study | |
| MKBMi254 | *B. microti* p*bspC*$_{\Delta SP}$-*phoA*$_{\Delta SP}$ | This study | |
| MKBMi255 | *B. microti* p*vceA-phoA*$_{\Delta SP}$ | This study | |
| MKBMi256 | *B. microti* Δ*virB9* p*vceA-phoA*$_{\Delta SP}$ | This study | |
| MKBMi257 | *B. microti* p*SP*$_{vceA}$-*phoA*$_{\Delta SP}$ | This study | |

| | | |
|---|---|---|
| MKBMi258 | *B. microti* p*vceA*$_{\Delta SP}$-*phoA*$_{\Delta SP}$ | This study |
| MKBMi229 | *B. microti* p*p17-phoA*$_{\Delta SP}$ | This study |
| MKBMi230 | *B. microti* p*ttpA-phoA*$_{\Delta SP}$ | This study |
| MKBMi231 | *B. microti* p*eipA-phoA*$_{\Delta SP}$ | This study |
| MKBMi148 | *B. microti* p*bspC-hibit* | This study |
| MKBMi171 | *B. microti* p*bpe123-hibit* | This study |
| MKBMi181 | *B. microti* p*vceA-hibit* | This study |
| MKBMi224 | *B. microti* p*p17-hibit* | This study |

Table S2: Plasmids used in this study.

| **Plasmid** | **Description** | **Reference** |
|---|---|---|
| pNPTS138 | Cloning suicide vector encoding *sacB*, $Kan^R$ | Kind gift of U. Jenal, Biozentrum of the University of Basel (CH) |
| pNPTS138_Δ*virB9*$_{BMi}$ | Suicide vector for in-frame deletion of *virB9* from *B. microti* | This study |
| pJS140 | pBBR-derivative with pTetAR2, dsRed under $P_{aphT}$, $Cm^R$ | (Ketterer *et al.* 2024) |
| p*phoA*$_{\Delta SP}$ | Derivative of pJS140, aTc-inducible expression of Flag-PhoA$_{\Delta SP}$ | This study |
| p*SP*$_{phoA}$-*phoA*$_{\Delta SP}$ | Derivative of p*phoA*$_{\Delta SP}$, aTc-inducible expression SP$_{PhoA}$- Flag-PhoA$_{\Delta SP}$ | This study |
| p*bspC*-*phoA*$_{\Delta SP}$ | Derivative of p*phoA*$_{\Delta SP}$, aTc-inducible expression BspC-Flag-PhoA$_{\Delta SP}$ | This study |
| p*SP*$_{bspC}$-*phoA*$_{\Delta SP}$ | Derivative of p*phoA*$_{\Delta SP}$, aTc-inducible expression SP$_{BspC}$-Flag-PhoA$_{\Delta SP}$ | This study |
| p*ha.bspC*$_{\Delta SP}$-*phoA*$_{\Delta SP}$ | Derivative of p*phoA*$_{\Delta SP}$, aTc-inducible expression BspC$_{\Delta SP}$-Flag-PhoA$_{\Delta SP}$ | |
| p*vceA*-*phoA*$_{\Delta SP}$ | Derivative of p*phoA*$_{\Delta SP}$, aTc-inducible expression VceA-Flag-PhoA$_{\Delta SP}$ | This study |
| p*SP*$_{vceA}$-*phoA*$_{\Delta SP}$ | Derivative of p*phoA*$_{\Delta SP}$, aTc-inducible expression SP$_{VceA}$-Flag-PhoA$_{\Delta SP}$ | This study |
| p*vceA*$_{\Delta SP}$ -*phoA*$_{\Delta SP}$ | Derivative of p*phoA*$_{\Delta SP}$, aTc-inducible expression VceA$_{\Delta SP}$-Flag-PhoA$_{\Delta SP}$ | This study |
| p*bspD*-*phoA*$_{\Delta SP}$ | Derivative of p*phoA*$_{\Delta SP}$, aTc-inducible expression BspD-Flag-PhoA$_{\Delta SP}$ | This study |
| p*eipA*-*phoA*$_{\Delta SP}$ | Derivative of p*phoA*$_{\Delta SP}$, aTc-inducible expression EipA-Flag-PhoA$_{\Delta SP}$ | This study |
| p*p17*-*phoA*$_{\Delta SP}$ | Derivative of p*phoA*$_{\Delta SP}$, aTc-inducible expression p17-Flag-PhoA$_{\Delta SP}$ | This study |
| p*ttpA*-*phoA*$_{\Delta SP}$ | Derivative of p*phoA*$_{\Delta SP}$, aTc-inducible expression TtpA-Flag-PhoA$_{\Delta SP}$ | This study |
| p*hibit* | Derivative of pJS140, aTc-inducible expression Flag-HiBiT for N-terminal fusions | This study |
| p*bspC*-*hibit* | Derivative of p*hibit*, aTc-inducible expression BspC-Flag-HiBiT | This study |
| p*ha.bspC*$_{\Delta SP}$-*hibit* | Derivative of p*hibit*, aTc-inducible expression HA.BspC$_{\Delta SP}$-Flaf-HiBiT | This study |
| p*vceA*-*hibit* | Derivative of p*hibit*, aTc-inducible expression VceA-Flag-HiBiT | This study |
| p*vceA*$_{\Delta SP}$-*hibit* | Derivative of p*hibit*, aTc-inducible expression VceA$_{\Delta SP}$-Flag-HiBiT | This study |
| p*bspL*-*hibit* | Derivative of p*hibit*, aTc-inducible expression BspL-Flag-HiBiT | This study |
| p*bpe123*-*hibit* | Derivative of p*hibit*, aTc-inducible expression BPE123-Flag-HiBiT | This study |
| p*p17*-*hibit* | Derivative of p*hibit*, aTc-inducible expression p17-Flag-HiBiT | This study |
| p*eipA*-*hibit* | Derivative of p*hibit*, aTc-inducible expression EipA-Flag-HiBiT | This study |

Table S3: Primers used in this study.

| Primer | Sequence | Purpose |
|---|---|---|
| prMK197 | TTGAAGCCGGCTGGCGCCAAGCTTCTCTGCAGTCCCGTCAACAAAAGCG | FW homology region 1 for Δ*virB9* from *B. abortus* 2308 |
| prMK198 | CCCGGGCGATTTGATGACCGCAAGCAGGAATCTTTTC | RV homology region 1 for Δ*virB9* from *B. abortus* 2308 |
| prMK199 | AAAAGATTCCTGCTTGCGGTCATCAAATCGCCCGG | FW homology region 2 for Δ*virB9* from *B. abortus* 2308 |
| prMK200 | AAGGCCTTGACTAGAGGGTCGACCTCGCTCGCAGAACACTTC | RV homology region 2 for Δ*virB9* from *B. abortus* 2308 |
| prMK201 | GTTGCGTGTGACCACG | Sequencing primer inside homology region 1 Δ*virB9* |
| prMK202 | CTTGCGGATCGTCACC | Sequencing primer Δ*virB9* |
| prMK203 | CGGTTTTGGTCGTACTACG | Sequencing primer Δ*virB9* |
| prMK117 | GAAAAGTGAAGAGCTCATGGATTATAAAGATCATGACGGTAGCGGTCGGACACCAGAAATGCC | FW Flag-PhoA without SP from *Escherichia coli* str. K12 substr. MG1655 for p*phoA*$_{\Delta SP}$ |
| prMK118 | CAATAACTGCCTTAAAAGCTTTTATTTCAGCCCCAGAGC | RV Flag-PhoA without SP from *Escherichia coli* str. K12 substr. MG1655 for p*phoA*$_{\Delta SP}$ |
| prMK123 | GATAGAGTTATTTTACCACTCCCTATCAGTGATAGAGAAAAGTGAAGAGCTCATGAAACAAAGCACTATTGCAC | FW SP$_{PhoA}$ from *Escherichia coli* str. K12 substr. MG1655 for p*SP*$_{phoA}$-*phoA*$_{\Delta SP}$ |
| prMK124 | GAACAGGCATTTCTGGTGTCCGACCGCTACCGTCATGATCTTTATAATCACCGCTACCGGCTTTTGTCACAGGGG | RV SP$_{PhoA}$ from *Escherichia coli* str. K12 substr. MG1655 for p*SP*$_{phoA}$-*phoA*$_{\Delta SP}$ |
| prMK151 | GATAGAGTTATTTTACCACTCCCTATCAGTGATAGAGAAAAGTGAAGAGCTCATGAAATCGACCAAGATCATACTTTC | FW *bspC* from *B. abortus* 2308 for p*bspC*-*phoA*$_{\Delta SP}$ and p*bspC*-*hibit* |
| prMK152 | AACAGGCATTTCTGGTGTCCGACCGCTACCGTCATGATCTTTATAATCACCGCTACCCTTGCGCACGATTTCTATG | RV *bspC* from *B. abortus* 2308 for p*bspC*-*phoA*$_{\Delta SP}$ and p*ha.bspC*$_{\Delta SP}$-*phoA*$_{\Delta SP}$ |
| prMK149 | GATAGAGTTATTTTACCACTCCCTATCAGTGATAGAGAAAAGTGAAGAGCTCATGAAATCGACCAAGATCATACTTTC | FW *SP*$_{bspC}$ from *B. abortus* 2308 for p*SP*$_{bspC}$-*phoA*$_{\Delta SP}$ |
| prMK150 | AACAGGCATTTCTGGTGTCCGACCGCTACCGTCATGATCTTTATAATCACCGCTACCGGCATCAGCTTGCGC | RV *SP*$_{bspC}$ from *B. abortus* 2308 for p*SP*$_{bspC}$-*phoA*$_{\Delta SP}$ |
| prMK210 | CACTCCCTATCAGTGATAGAGAAAAGTGAAGAGCTCATGTACCCATACGATGTTCCAGATTACGCTGTGCCAAAGCGGACAAAG | FW *ha.bspC*$_{\Delta SP}$ from *B. abortus* 2308 for p*ha.bspC*$_{\Delta SP}$-*hibit* and p*ha.bspC*$_{\Delta SP}$-*phoA*$_{\Delta SP}$ |
| prMK163 | GATAGAGTTATTTTACCACTCCCTATCAGTGATAGAGAAAAGTGAAGAGCTCATGAAAATCATCATCACGGCAGC | FW *vceA* from *B. abortus* 2308 for p*vceA*-*phoA*$_{\Delta SP,}$ p*SP*$_{vceA}$-*phoA*$_{\Delta SP}$, and p*vceA*-*hibit* |
| prMK164 | AACAGGCATTTCTGGTGTCCGACCGCTACCGTCATGATCTTTATAATCACCGCTACCGTTCTTGGGCGCGTGGC | RV *vceA* from *B. abortus* 2308 for p*vceA*-*phoA*$_{\Delta SP}$ and p*vceA*$_{\Delta SP}$ -*phoA*$_{\Delta SP}$ |
| prMK165 | AACAGGCATTTCTGGTGTCCGACCGCTACCGTCATGATCTTTATAATCACCGCTACCGGCCATTGCGCCCGTG | RV *SP*$_{vceA}$ from *B. abortus* 2308 for p*SP*$_{vceA}$-*phoA*$_{\Delta SP}$ |
| prMK166 | GATAGAGTTATTTTACCACTCCCTATCAGTGATAGAGAAAAGTGAAGAGCTCATGCAAAAAACCCAATGCGATGCAAAG | FW *vceA*$_{\Delta SP}$ from *B. abortus* 2308 for p*vceA*$_{\Delta SP}$-*phoA*$_{\Delta SP}$, and p*vceA*$_{\Delta SP}$-*hibit* |
| prMK131 | GATAGAGTTATTTTACCACTCCCTATCAGTGATAGAGAAAAGTGAAGAGCTCATGATTCAATCCGCTCTCTTTTTG | FW *bspD* from *B. abortus* 2308 for p*bspD*-*phoA*$_{\Delta SP}$ |
| prMK121 | CTACCGTCATGATCTTTATAATCACCGCTACCTTGCATGTCGCGGATGC | RV *bspD* from *B. abortus* 2308 for p*bspD*-*phoA*$_{\Delta SP}$ |

| | | |
|---|---|---|
| prMK156 | GATAGAGTTATTTTACCACTCCCTATCAG TGATAGAGAAAAGTGAAGAGCTCATGAAT CGATTTTTGAAGATCACTATTC | FW *bspL* from *B. abortus* 2308 for p*bspL-hibit* |
| prMK114 | GATAGAGAAAAGTGAAGAGCTCATGAGC TTGTTGCTGGCTAAC | FW *bpe123* from *B. abortus* 2308 for p*bpe123-hibit* |
| prMK216 | CCTTGTAATCGATGTCATGATCTTTATAAT CACCGTCATGGTCTTTGTAGTCTGCCTGT CCCGCCAG | RV *bpe123* from *B. abortus* 2308 for p*bpe123-hibit* |
| prMK214 | CACTCCCTATCAGTGATAGAGAAAAGTGA AGAGCTCATGGCCATACCATCCCTG | FW *eipA* from *B. abortus* 2308 for p*eipA-phoA*$_{\Delta SP}$, and p*eipA-hibit* |
| prMK278 | CTACCGTCATGATCTTTATAATCACCGCT ACCGAACGGGTTCCAGGTCG | RV *eipA* from *B. abortus* 2308 for p*eipA-phoA*$_{\Delta SP}$ |
| prMK268 | GATAGAGTTATTTTACCACTCCCTATCAG TGATAGAGAAAAGTGAAGAGCTCATGAAC CAAAGCTGTCCG | FW *p17* from *B. abortus* 2308 for p*p17-phoA*$_{\Delta SP}$, and p*p17-hibit* |
| prMK276 | CTACCGTCATGATCTTTATAATCACCGCT ACCGACAAGCGCGGCGAT | RV *p17* from *B. abortus* 2308 for p*p17-phoA*$_{\Delta SP}$ |
| prMK272 | GATAGAGTTATTTTACCACTCCCTATCAG TGATAGAGAAAAGTGAAGAGCTCATGCG GCTAGGATCGAAAC | FW *ttpA* from *B. abortus* 2308 for p*ttpA-phoA*$_{\Delta SP}$ |
| prMK277 | CTACCGTCATGATCTTTATAATCACCGCT ACCATCGGTTGTCGGCTTTTC | RV *ttpA* from *B. abortus* 2308 for p*ttpA-phoA*$_{\Delta SP}$ |
| prMK206 | GAAAAGTGAAGAGCTCATGGACTACAAA GACCATGACGGTG | FW *hibit* for p*hibit* |
| prMK207 | CAATAACTGCCTTAAAAGCTTTTAGCTAAT CTTCTTGAACAGCCG | RV *hibit* for p*hibit* |
| prMK208 | CCTTGTAATCGATGTCATGATCTTTATAAT CACCGTCATGGTCTTTGTAGTCCTTGCGC ACGATTTCTATG | RV *bspC* from *B. abortus* 2308 for p*bspC-hibit* and p*ha.bspC*$_{\Delta SP}$*-hibit* |
| prMK252 | CTTTATAATCACCGTCATGGTCTTTGTAG TCGTTCTTGGGCGCGTGGC | RV *vceA* from *B. abortus* 2308 for p*vceA-hibit* and p*vceA*$_{\Delta SP}$*-hibit* |
| prMK251 | CTTTATAATCACCGTCATGGTCTTTGTAG TCGTTGGCCGTGCAGAAATG | RV *bspL* from *B. abortus* 2308 for p*bspL-hibit* |
| prMK269 | CTTTATAATCACCGTCATGGTCTTTGTAG TCGACAAGCGCGGCGAT | RV *p17* from *B. abortus* 2308 for p*p17-hibit* |
| prMK215 | CCTTGTAATCGATGTCATGATCTTTATAAT CACCGTCATGGTCTTTGTAGTCGAACGG GTTCCAGGTCG | RV *eipA* from *B. abortus* 2308 for p*eipA-hibit* |

Table S4: *In silico* analysis of *B. abortus* effector proteins with T4SS-dependent translocation predicted signal peptides and N-proximal transmembrane domains for a subset of putative effector proteins.

| **Effector** | **ORF**[1] | **Size [kDa]** | **SignalP6.0 [Likelihood]** | | **Phobius** | | **DeepTMHMM** | | **Translocation/Secretion mechanism** |
|---|---|---|---|---|---|---|---|---|---|
| | | | Sec/SPI (Signal Peptide)[2] | Sec/SPII (Lipoprotein Signal Peptide)[2] | Sec/SPI (Signal Peptide)[2] | N-proximal Transmembrane Domain[2,3] | N-proximal Transmembrane Domain[2,3] | Signal Peptide | |
| VceA | BAB1_1652 | 11.3 | Yes | - | Yes | - | - | Yes | T4SS-dependent translocation (De Jong *et al.* 2008) |
| VceC | BAB1_1058 | 44.9 | - | - | - | Yes | Yes | - | T4SS-dependent translocation (De Jong *et al.* 2008) |
| BPE005 | BAB1_2005 | 16.8 | - | - | - | - | - | - | T4SS-dependent translocation (Marchesini *et al.* 2011) |
| BPE043 | BAB1_1043 | 166.4 | - | - | - | - | - | - | T4SS-dependent translocation (Marchesini *et al.* 2011) |
| BPE123 | BAB2_0123 | 16.7 | - | - | Yes | - | Yes | - | T4SS-dependent translocation, periplasmic intermediate (Marchesini *et al.* 2011; Del Giudice *et al.* 2016) |
| BPE275 | BAB1_1275 | 27.7 | - | - | - | Yes | Yes | - | T4SS-dependent translocation (Marchesini *et al.* 2011) |
| SepA | BAB1_1492 | 21.0 | - | - | - | - | - | - | T4SS-dependent translocation, periplasmic intermediate (Döhmer *et al.* 2014) |
| BtpA/Btp1 | BAB1_0279 | 28.0 | - | - | - | - | - | - | T4SS-dependent translocation (Salcedo *et al.* 2013) |
| BtpB/Btp2 | BAB1_0756 | 31.8 | - | - | - | - | - | - | T4SS-dependent translocation (Salcedo *et al.* 2013) |
| BspA | BAB1_0678 | 21.7 | - | - | - | Yes | Yes | - | T4SS-dependent translocation (Myeni *et al.* 2013) |
| BspB | BAB1_0712 | 20.5 | - | - | Yes | - | Yes | - | T4SS-dependent translocation (Myeni *et al.* 2013) |
| BspC | BAB1_0847 | 14.8 | Yes | - | Yes | - | - | Yes | T4SS-dependent translocation (Myeni *et al.* 2013) |
| BspE | BAB1_1675 | 12.7 | - | - | - | - | - | - | T4SS-dependent translocation (Myeni *et al.* 2013, 2025) |
| BspF | BAB1_1948 | 48.8 | - | - | - | - | - | - | T4SS-dependent translocation (Myeni *et al.* 2013) |
| BspL | BAB1_1533 | 18.8 | Yes | - | Yes | - | - | Yes | T4SS-dependent translocation (Luizet *et al.* 2021) |
| NyxA | BAB1_0296 | 14.5 | - | - | - | - | - | - | T4SS-dependent translocation (Louche *et al.* 2023) |
| NpeA | BAB2_0195 | 24.4 | - | Yes | Yes | - | - | Yes | T4SS-dependent translocation, periplasmic intermediate (Giménez *et al.* 2024) |
| RS14565 | BAB2_0295 | 33.4 | - | - | - | - | - | - | T4SS-dependent secretion, translocation into host cell (Yin *et al.* 2024) |
| RS15060 | BAB2_1002 | 35.8 | - | - | - | - | - | - | T4SS-dependent secretion, translocation into host cell (Yin *et al.* 2024) |
| RS10635 | BAB2_0074 | 35.7 | - | - | Yes | - | Yes | - | T4SS-dependent secretion, translocation into host cell (Yin *et al.* 2024) |
| Rhg1 | BAB1_0478 | 33.2 | - | - | - | Yes | Yes | - | T4SS-dependent translocation (Cabello *et al.* 2024) |

[1] open reading frame in *B. abortus* 2308 genome
[2] Yes = Probability ≥0.9
[3] within N-proximal 50 aa

Table S5: *In silico* analysis of *B. abortus* secreted and translocated proteins with unresolved translocation or secretion mechanism predicted signal peptides and N-proximal transmembrane domains for a subset of putative effector proteins.

| Effector | ORF[1] | Size [kDa] | SignalP6.0 [Likelihood] | | Phobius | | DeepTMHMM | | Translocation/Secretion mechanism |
|---|---|---|---|---|---|---|---|---|---|
| | | | Sec/SPI (Signal Peptide) [2] | Sec/SPII (Lipoprotein Signal Peptide) [2] | Sec/SPI (Signal Peptide)[2] | N-proximal Transmembrane Domain[2,3] | N-proximal Transmembrane Domain[2,3] | Signal Peptide | |
| RicA | BAB1_1279 | 18.6 | - | - | - | - | - | - | T4SS-dependent translocation, T4SS-independent secretion (De Barsy *et al.* 2011) |
| BPE159 | BAB2_0159 | 18.5 | - | - | - | Yes | Yes | - | T4SS-independent translocation (Marchesini *et al.* 2011) |
| BPE865/BspI | BAB1_1865 | 24.9 | - | - | - | - | - | - | Translocated, unclear mechanism (Marchesini *et al.* 2011; Myeni *et al.* 2013) |
| BspD | BAB1_1611 | 34.9 | - | - | - | Yes | Yes | - | Translocated, unclear mechanism (Myeni *et al.* 2013) |
| BspG | BAB1_0277 | 23 | - | - | - | - | - | - | T4SS-independent translocation (Myeni *et al.* 2013, 2025) |
| BspH | BAB1_1864 | 53.7 | - | - | - | - | - | - | Translocated, unclear mechanism (Myeni *et al.* 2013) |
| BspJ | BAB2_0119 | 19.6 | - | - | - | - | - | - | Translocated, unclear mechanism (Myeni *et al.* 2013; Ma *et al.* 2020) |
| BspK | BAB2_0541 | 14.3 | - | - | - | Yes | Yes | - | T4SS-independent translocation (Myeni *et al.* 2013) |
| NyxB | BAB1_1101 | 15.0 | - | - | - | - | - | - | T4SS-independent secretion (Louche *et al.* 2023) |
| CstA | BAB2_0856 | 51.7 | - | - | - | - | - | - | T4SS-independent secretion (De Barsy *et al.* 2012) |
| SodC | BAB2_0535 | 18.1 | Yes | - | Yes | - | - | Yes | T4SS-independent secretion, periplasmic protein (Beck, Tabatabai and Mayfield 1990; Bricker *et al.* 1990; Liu *et al.* 2018) |

[1] open reading frame in *B. abortus* 2308 genome
[2] Yes = Probability ≥0.9
[3] within N-proximal 50 aa

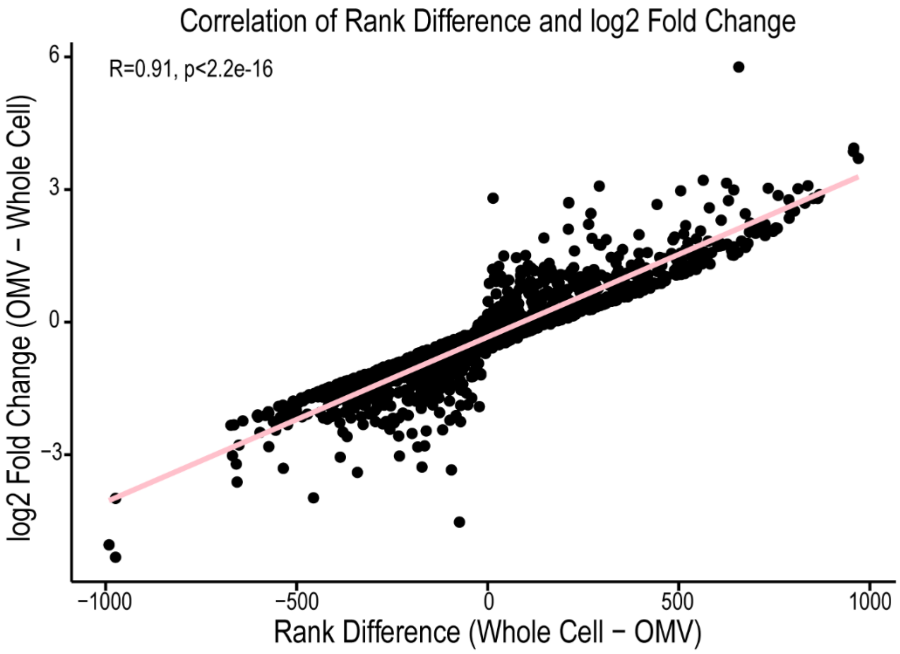

**Figure S1:** Correlation of rank difference between proteins quantified in whole cells and OMVs and log2 fold change of proteins quantified in OMVs vs whole cells.

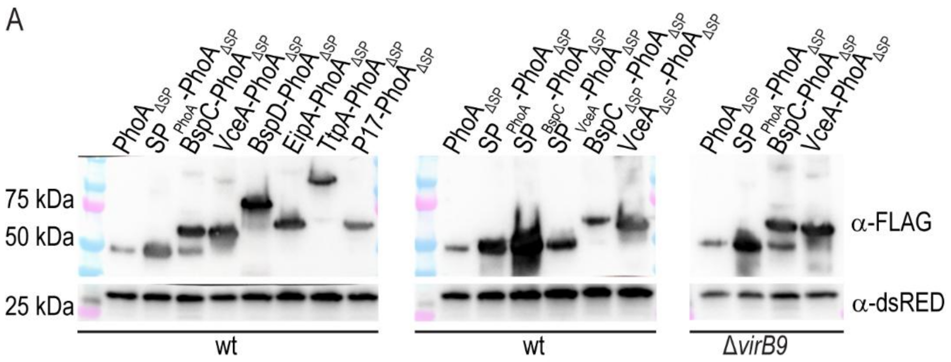

**Figure S2:** Western Blots to control expression of PhoA-fusion proteins. **(A)** Equal amounts of bacterial pellet were loaded onto an SDS-PAGE and visualized using Western Blotting. Shown is one representative replicate of two.

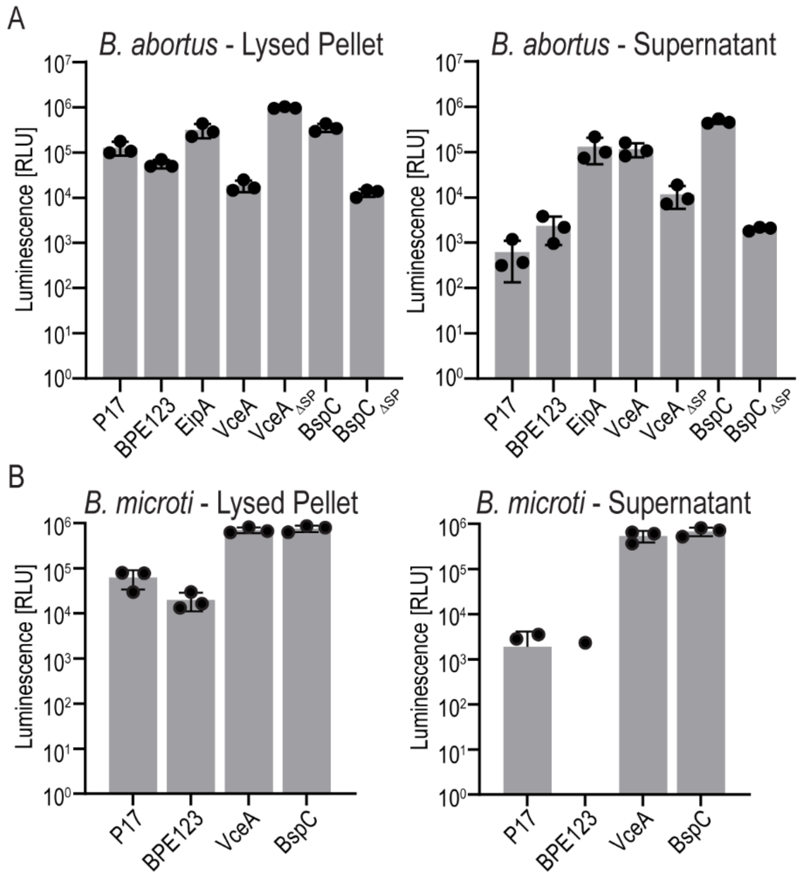

**Figure S3:** RAW luminescence measurements corresponding to Figure 2A and D. RLU = Relative Light Units. Log-scale. Background substracted. n=3. **(A)** Luminescence measurements from lysed *B. abortus* expressing HiBiT-fusion proteins (left) and bacteria-free supernatant (right) supplemented with LgBiT. **(B)** Luminescence measurements from lysed *B. microti* expressing HiBiT-fusion proteins (left) and bacteria-free supernatant (right) supplemented with LgBiT. In the supernatant fraction some values for P17 and BPE123 were negative after background subtraction and are therefore not blotted.

**Data S1_SupplementaryData**. For each identified protein, the orthogroup was generated using OrthoFinder version 2.5.5 to match *Brucella* proteins between species to compare published proteomes at the orthogroup level.

**Data S2_SupplementaryData.** Lists identified ions and peptides, as well as proteins from three replicates and associated search engine scores. Also listed are proteins exclusively found in the whole cell or OMV-fractions, named “WCExcl” and “OMVExcl”, respectively.

**Data S3_SupplementaryData.** Lists the quantified proteins from 3 replicates (“AnnotatedQuantData”) and their scores. Also listed are proteins exclusively found in the whole cell or OMV-fractions, named “WCExcl” and “OMVExcl”, respectively.

**Data S4_SupplementaryData.** Lists all proteins that were quantified at least once in OMVs and whole-cell samples ranked by abundance and log2 fold change. “Marker Rank Difference” lists the rank difference and log2 fold change of all marker proteins depicted in Fig. 1A and B. “T4SS Rank and FC” lists the rank difference and log2 fold change of T4SS effectors depicted in Fig. 1C.

**Data S5_SupplementaryData.** Shows in how many of the three replicates the listed effectors were identified and quantified.

**Data S6_SupplementaryData.** Lists the identified effectors and indicates their presence or absence in OMV proteomes of *B. melitensis* (Avila-Calderón *et al.* 2020), *B. abortus* (Araiza-Villanueva *et al.* 2019), and *B. suis* (Socorro Ruiz-Palma *et al.* 2021), as well as if the protein was quantified in the *B. abortus* OMVs and if it was enriched or depleted.
